# Nuclear poly-glutamine aggregates rupture the nuclear envelope and hinder its repair

**DOI:** 10.1101/2022.11.09.515785

**Authors:** Giel Korsten, Robin Pelle, Baukje Hoogenberg, Harm H. Kampinga, Lukas C. Kapitein

## Abstract

Huntington’s disease (HD) is caused by a poly-glutamine expansion of the huntingtin protein, resulting in the formation of poly-glutamine aggregates. The mechanisms of toxicity that result in the complex HD pathology remain only partially understood. Here we show that nuclear polyglutamine aggregates deform the nuclear envelope (NE) and induce NE ruptures that are often repaired incompletely. These ruptures coincide with deformations of the nuclear lamina and lead to lamina scar formation. Expansion microscopy enabled resolving the ultrastructure of nuclear aggregates and revealed polyglutamine fibrils sticking into the cytosol at rupture sites, suggesting a mechanism for incomplete repair. These findings implicate nuclear polyQ aggregate-induced loss of NE integrity as a potential contributing factor to Huntington’s disease and other polyglutamine diseases.

**One-sentence summary:** Aggregates associated with Huntington’s Disease induce ruptures of the nuclear envelop that compromise its barrier function

## Main Text

Huntington’s disease (HD) is a debilitating neurodegenerative disease caused by a CAG repeat expansion in exon-1 of the huntingtin gene (*1*), resulting in the expression of polyglutamine (polyQ) proteins that are prone to aggregate. A unified understanding of HD pathology is lacking, but polyQ aggregation has been associated with a wide range of cellular defects, including reduced protein quality control and transcriptional deregulation (*2–6*), as well as a compromised barrier function of the nuclear envelope (NE) (*7, 8*). While the latter has primarily been attributed to impaired nucleocytoplasmic shuttling due to sequestration of shuttling factors (*7, 8*), cells with nuclear polyQ aggregates also display abnormalities in their nuclear lamina (*7–9*), an intranuclear cytoskeletal scaffold that serves to protect the NE from rupturing (*10*). This suggests that ruptures of the NE might contribute to impaired nuclear barrier function in HD. Consistently, recent work has revealed that polyQ aggregates can directly disrupt organelle membranes (*11, 12*) and interact with the NE (*9, 13–15*). However, whether such interactions result in polyQ aggregate-induced ruptures that compromise the barrier function of the NE has remained unexplored.

To study the effect of polyQ aggregates on NE integrity, we expressed various forms of huntingtin exon1 (*16*) in cells stably expressing RFP with a nuclear localization signal (U2OS-RFP-NLS; Fig. 1A) (*17*). As expected, an expanded form of huntingtin targeted to the nucleus (polyQ74-NLS) exclusively formed nuclear aggregates, while non-expanded (polyQ23-NLS) or non-targeted (polyQ74) huntingtin formed no, or only cytoplasmic inclusions in this cell line, respectively (Fig. 1B). We then performed long-term (8 hours) live-cell imaging of these cells and found that cells with nuclear aggregates frequently showed loss of NE integrity (32.9±8.6%, n = 325 cells; Fig. 1, C-E; Movie S1), demonstrated by a rapid loss of RFP from the nucleus. Single cells often showed multiple rounds of rupture (1.8±1.3 ruptures per cell, n = 108 cells) and repair, indicated by the reaccumulation of RFP in the nucleus (fig. S1, A-D). In contrast, expression of either cytosolic or non-expanded polyQ protein only resulted in a minor increase in NE ruptures (7.7±8.3% and 7.2±1.9%, n = 165 and 506 cells) compared to control (2.0±0.8%, n = 1069 cells; Fig. 1E). Ruptures were sometimes, but not always, preceded by NE blebbing events (Fig. 1, C, F and G; fig. S1B), similar to the NE herniations shown to arise during constricted migration or after lamin depletion (*18–20*). While NE blebbing also occurred at higher frequency in cells expressing non-expanded or cytosolic aggregates (12.6±0.4% and 24.0±6.8%), they were most frequent in cells with nuclear aggregates (39.6±7.0%; Fig. 1F). These findings demonstrate that nuclear polyQ aggregates induce NE blebbing and rupture, reflecting a compromised barrier function of the NE.

**Fig. 1.**
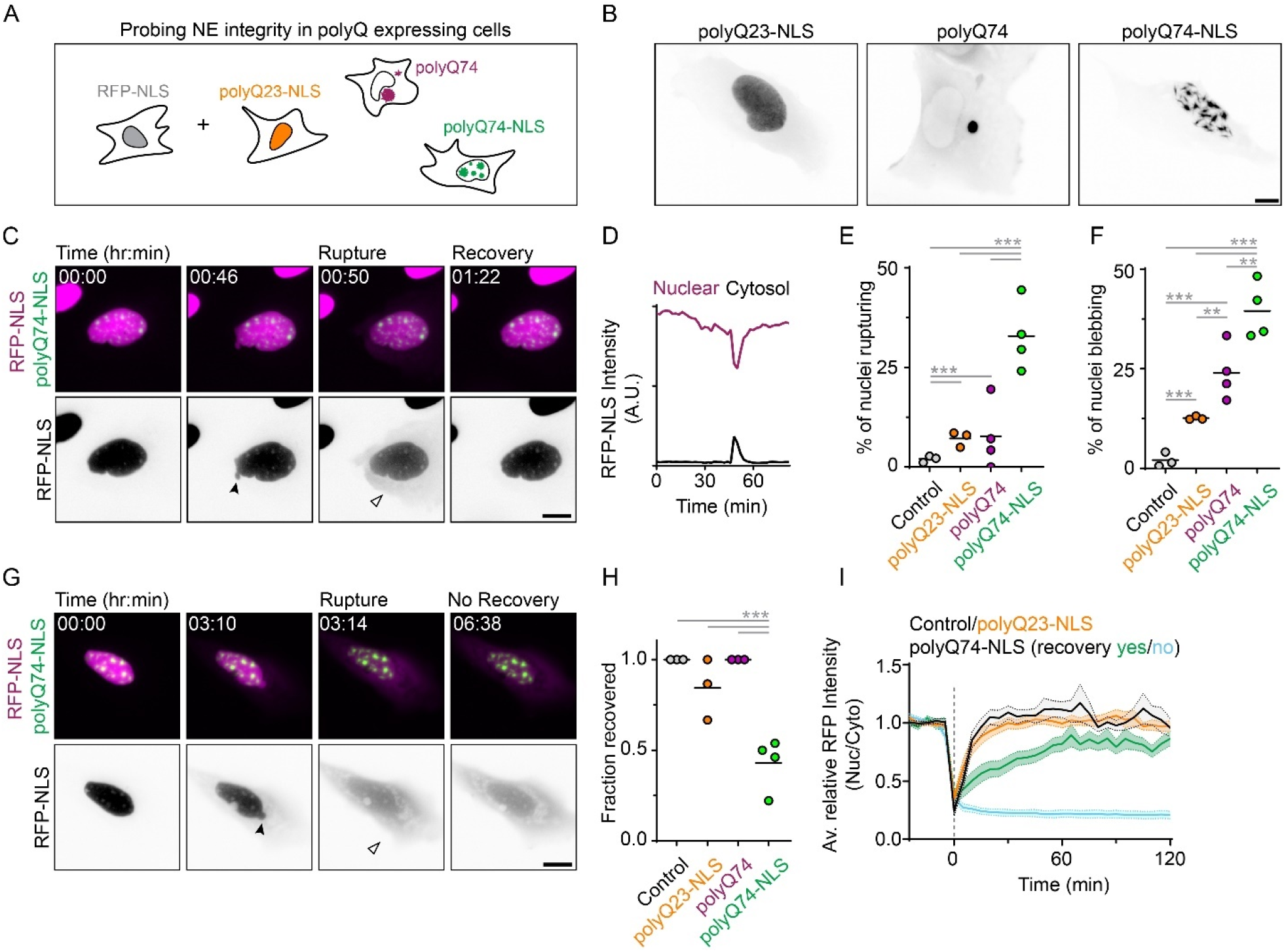
Nuclear aggregates induce NE rupture and impair recovery. (**A**) Schematic representation of experimental design probing NE integrity in U2OS-RFP-NLS (grey) cells expressing nuclear (polyQ74-NLS), non-expanded (polyQ23-NLS) or non-targeted (polyQ74) polyQ protein. (**B**) Representative images of localization and aggregate formation in cells expressing various polyQ constructs. (**C**) Timelapse images of NE rupture in representative U2OS-RFP-NLS (magenta) cell expressing polyQ74-NLS (green), resulting in transient loss of nuclear RFP-NLS enrichment. Solid arrow head marks nuclear bleb, open arrow head mark cytosolic RFP-NLS. (**D**) Nuclear (magenta curve) and cytosolic (black curve) RFP-NLS intensity of cell in (C). (**E and F**) Percentage of cells showing NE rupture (E) and NE blebbing (F) in U2OS-RFP-NLS control cells and cells expressing polyQ23-NLS, polyQ74cyto and polyQ74-NLS (n=1069, 506, 163, 325; N=3,3,4,4). (**G**) Timelapse images of NE rupture in representative U2OS-RFP-NLS cells expressing polyQ74-NLS, showing permanent loss of RFP-NLS enrichment after NE rupture. (**H**) Fraction of U2OS-RFP-NLS cells that recovered nuclear enrichment of RFP-NLS signal after NE rupture (n=21, 37, 13, 103; N=3,3,4,4). (**I**) Recovery of normalized nuclear RFP-NLS enrichment after rupture (t=0) in control cells (black curve) and cells expressing polyQ23-NLS (orange curve) and polyQ74-NLS cells with (green curve) or without recovery (cyan curve; n=18, 21, 17, 31; N=3,3,4,4). Horizontal bars represent mean ± SEM. Scale bars are 10 μm (B) or 15 μm (C and G). *p<0.05, ***p<0.001.

While NE ruptures are typically quickly repaired, as revealed by the reaccumulation of RFP-NLS (*17, 19–21*), we noticed multiple instances of permanent loss of NE integrity after rupture in cells with nuclear aggregates (Fig. 1G; Movie S2; fig. S1, A-C). To determine whether this impaired recovery was specific for cells with nuclear aggregates, we scored the fraction of cells that recovered after NE rupture and determined their recovery dynamics. Indeed, ruptures in cells with nuclear aggregates recovered less often (43±14% recovery; Fig. 1H) and recovered slower than in control cells or in cells expressing non-expanded polyQ protein (Fig. 1I). These results suggest that nuclear aggregates interfere with NE resealing (*18*).

Since both NE ruptures and impaired recovery were specifically induced by nuclear aggregates, we hypothesized that these aggregates would locally deform and disrupt the nuclear lamina and NE. First, we determined whether ruptures indeed occurred close to nuclear aggregates. To this end, we co-expressed polyQ74-NLS with mCherry-tagged guanosine 3’,5’-monophosphate–adenosine3’,5’-monophosphate (cyclic GMP-AMP) synthase (cGAS), a DNA binding protein rapidly recruited to sites of NE rupture (*21, 22*). Indeed, ruptures occurred specifically at nuclear aggregates, as evidenced by accumulation of endogenous (fig. S2A) and mCherry-cGAS around aggregates (i.e. within 1.3±1 μm, n = 25 ruptures; Fig. 2, A-C; Movie S3; fig. S2C). In some cells mCherry-cGAS accumulated around multiple distinct aggregates at different times during imaging, suggesting that the ruptures previously observed in U2OS-RFP-NLS cells occurred at different sites (fig. S2B and C; fig. S1, A-D).

**Fig. 2.**
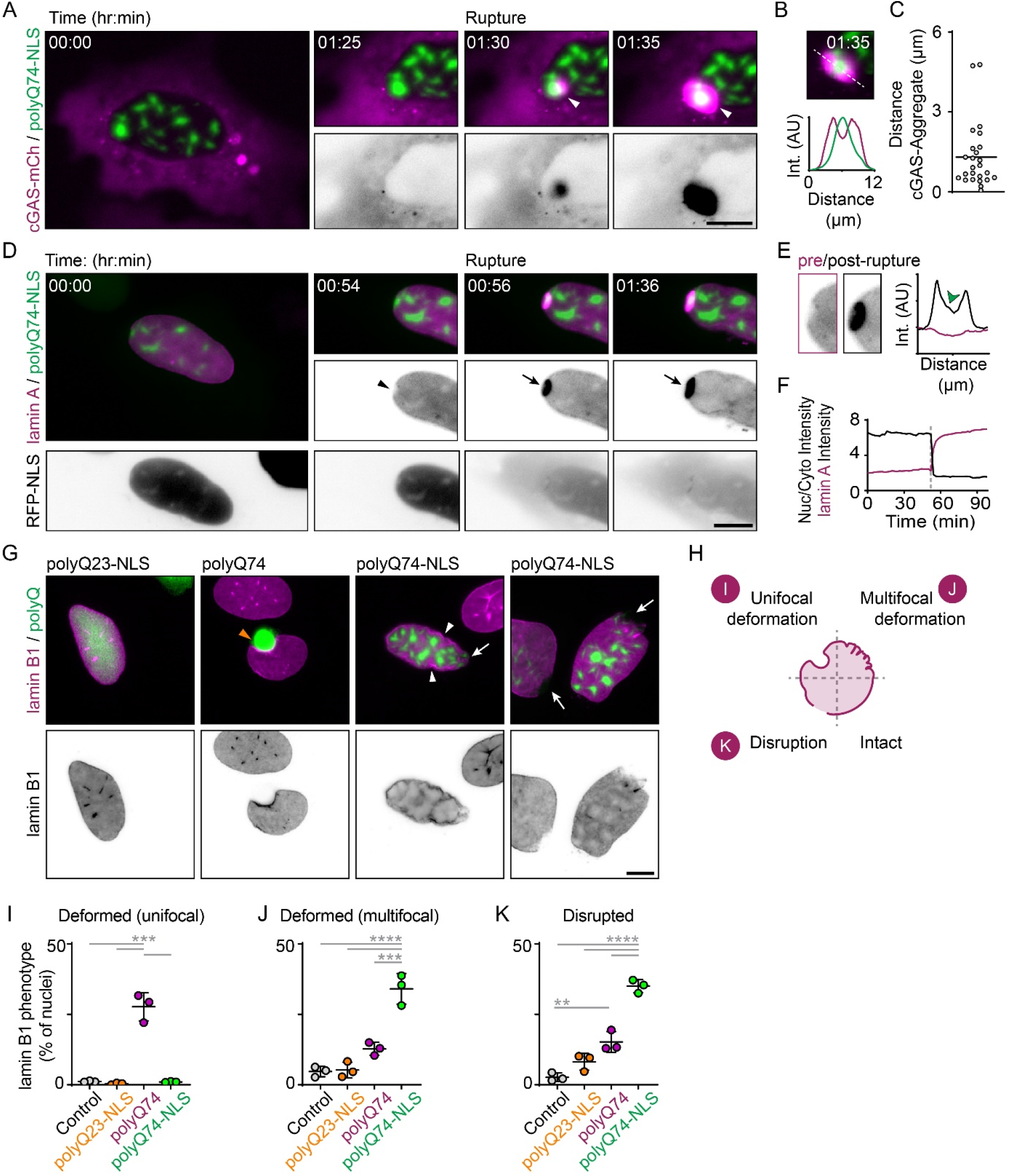
Nuclear aggregates induce local disruptions of the nuclear lamina and nuclear envelope. (**A**) Timelapse images of representative U2OS cell expressing cGAS (magenta) and polyQ74-NLS (green), showing nuclear cGAS entry (white arrowhead) around a nuclear aggregate. (**B**) Intensity profile of cGAS accumulation around aggregate shown in (A). **(C**) Distance between cGAS accumulation and polyQ74-NLS aggregate 0-5 min. after NE rupture (n=25; N=3). (**D**) Live-cell imaging of representative U2OS-RFP-NLS (grey) cells expressing polyQ74-NLS (green) and HaloTag-lamin A (magenta) showing local lamin A depletion (arrowhead) and scar formation post-rupture (black arrowhead). (**E**) Zoom and intensity profile of rupture site shown in (D) pre- and post-rupture. (**F**) Graph showing increase in lamin A intensity (magenta) at rupture site after loss of RFP-NLS nuclear enrichment (black) of cell shown in (D). (**G**) Representative U2OS cells expressing polyQ23-NLS, polyQ74-NLS or polyQ74cyto (green) immunostained for lamin B1 (magenta), showing lamin B1 disruption (white arrows), unifocal (orange arrowhead) and multifocal (white arrowheads) lamin B1 deformation. (**H**) Schematic representation of lamin B1 phenotypes shown in (I-K). (**I-K**) Percentage of U2OS control cells, or cells expressing polyQ23-NLS, polyQ74cyto or polyQ74-NLS that show unifocal deformation (I), multifocal deformation (J) or disruption (K) of lamin B1. PolyQ (A and D) and cGAS (A) signal was gamma adjusted (γ = 0.75). Horizontal bars represent mean ± SD (n=589, 480, 253, 289; N=3). All scale bars represent 10 μm.). **p<0.01, ***p<0.001, ****p<0.0001.

Next, we tested whether these sites of aggregate-induced rupture also display lamina deformations. We expressed nuclear aggregates and HaloTag-lamin A in our reporter cells and found that lamin A intensity was often reduced near nuclear aggregates, but rapidly accumulated at aggregates upon NE rupture (Fig. 2, D-F; Movie S4; fig. S3, A-D). This accumulation of lamin A partly resembles the “scar” formation following rupture that was previously reported and hypothesized to be locally protective (*19*). Following aggregate-induced rupture, however, lamin A accumulation seemed incomplete and disrupted by the presence of the aggregate (Fig. 2E). We also observed occasional accumulation of HaloTag-lamin A at nuclear aggregates prior to NE rupture, possibly reflecting the repair of a destabilizing nuclear lamina (fig. S3, E and F). To validate and quantify these findings for endogenous lamins, we used immunostaining and analyzed nuclear deformations in different conditions (Fig. 2G). While cytosolic aggregates often induced a single, large nuclear deformation (27.6±4.9%, n = 253 cells; Fig. 2, G-I), nuclear aggregates induced lamin B1 deformations throughout the nuclear envelope (34.1±5.4%, n = 289 cells; Fig. 2, G, H and J), reminiscent of lamina “wrinkling” associated with HD nuclei (*7, 9, 23*). Nuclear aggregates furthermore induced dissociation of parts of the lamin B1 meshwork, resulting in areas devoid of lamin B1 (35.0±2.4%, n = 289 cells; Fig. 2, G, H and K) that are likely prone to NE rupture (*19*). Interestingly, these disruptions were only rarely present in cells with cytosolic aggregates (12.8±2.3%, n = 253 cells; Fig. 2, G and K). Together, these findings suggest that nuclear aggregates locally deform and disrupt the nuclear lamina, resulting in frequent NE ruptures near aggregates.

To facilitate a nanoscale analysis of local deformations and disruptions near nuclear aggregates, we turned to tenfold robust expansion microscopy (TREx), which enables specific labelling of aggregates and lamina in combination with visualization of the NE membranes. Expanded cells displaying different degrees of aggregation revealed a striking improvement in resolution compared to confocal microscopy (Fig. 3A; Movie S5), allowing three-dimensional visualization and segmentation of individual nuclear aggregates. These aggregates had dense cores with individual fibrils protruding outward (Fig. 3, B and C), similar to polyQ fibrils protruding from cytosolic inclusions observed in EM (*11, 12*).

**Fig. 3.**
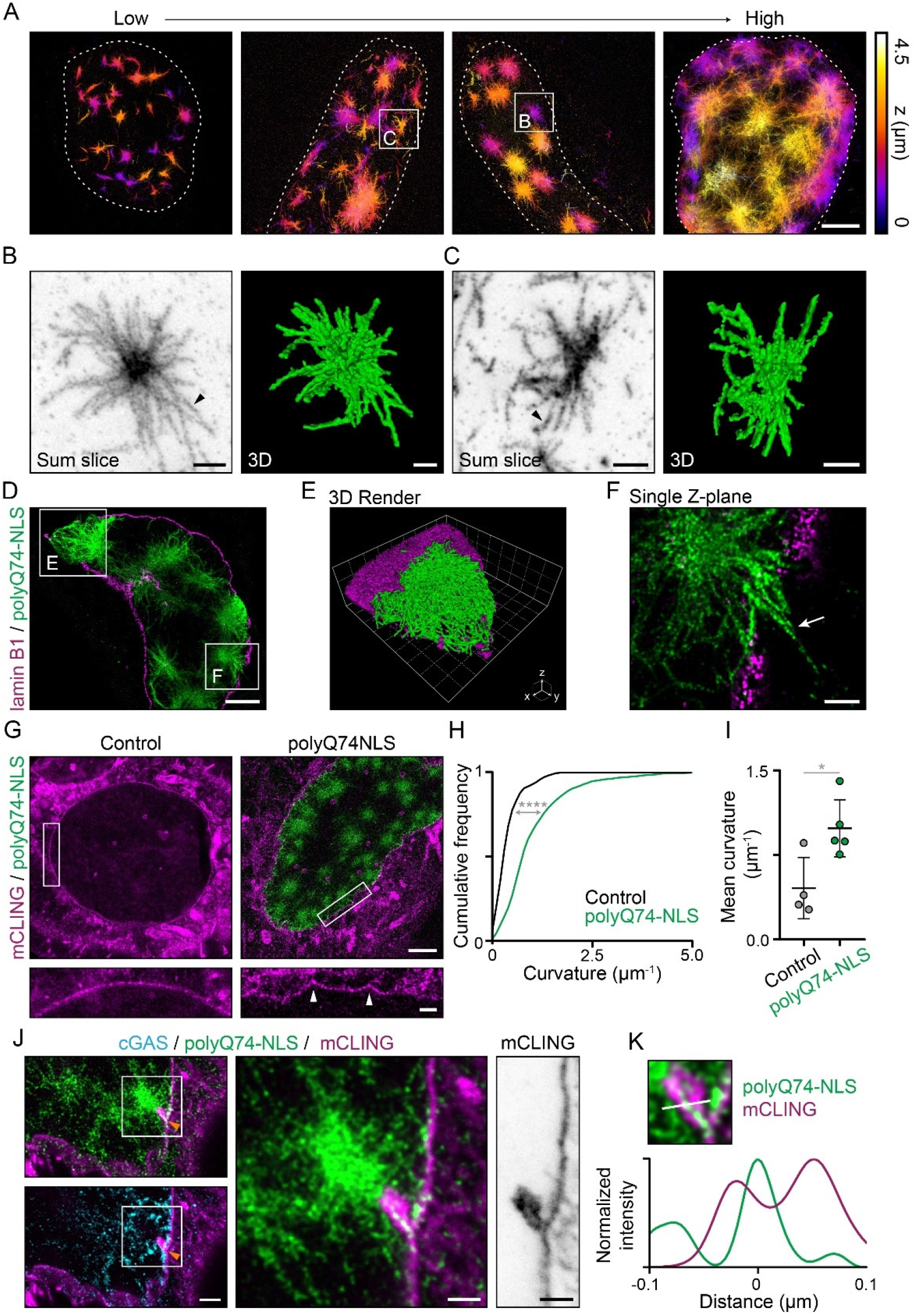
Ultrastructure of nuclear aggregates and nuclear envelope deformations visualized using TREx microscopy. (**A**) Depth-encoded color projection of representative expanded U2OSWT cells expressing polyQ74-NLS, showing various stages of aggregation progression (low to high). See also Movie S5. (**B-C**) Sum projection (inverted grey scale) and 3D volumetric render (green) of single aggregates shown in (A), showing distinct fibrillar structures protruding from a nuclear aggregate (arrowheads). (**D**) Representative image of expanded U2OSWT cell expressing polyQ74-NLS (green), stained for lamin B1 (magenta). (**E**) 3D volumetric render of lamin B1 disruption shown in (D). (**F**) Single z-plane of cell in (D). Representative event of polyQ74-NLS fibrils protruding through a lamin B1 disruption (white arrow). (**G**) Representative expanded U2OSWT control cell and cell expressing polyQ74-NLS, stained with total membrane stain (mCling; magenta). Zooms show a stretch of continuous nuclear membrane with high local curvatures (white arrowheads) in cells with nuclear aggregates. (**H-I**) Distribution of local membrane curvatures (H) and mean curvature per cell (I) (Control (black) and polyQ74-NLS (green); 4 and 5 cells). (**J**) Cell expressing polyQ74-NLS and cGAS (cyan), showing nuclear membrane deformation around a nuclear aggregate, indicated by cGAS accumulation (orange arrowheads). (**K**) Zoom and line profile of the rupture site in (J) showing invagination of NE membrane around aggregate fibril. Scale bars: 2.5 μm (A, D and G), 500 nm (B, C, F, G_zoom_ and J), 250 nm (J_zoom_). Voxel sides are 600 nm (grid of E). *p<0.05, ****p<0.0001.

Next, labelling of nuclear aggregates and endogenous lamin B1 allowed us to observe the structure of the lamin B1 meshwork using TREx (fig. S4, A-C). In control cells, lamin B1 was present as a continuous meshwork lining the nuclear membrane, resembling earlier results obtained using other super-resolution techniques (*24–26*) (fig. S4, A-C). Consistent with our earlier data (Fig. 2, G, J and K), cells with nuclear aggregates displayed strong lamin B1 abnormalities. Nuclear aggregates colocalized with large disruptions (fig. 3, D and E), as well as smaller disruptions where polyQ fibrils appeared to protrude through holes in the lamina meshwork (Fig. 3, D and F; fig. S4D). In contrast, the lamin B1 meshwork appeared intact near cytosolic aggregates, consistent with earlier reports (*11, 12*) (fig. S4E). In addition to the lamin B1 discontinuities found near nuclear aggregates, we noticed that the overall shape of the lamina was more irregular in cells with nuclear aggregates than in control cells. Quantification of the local lamina curvatures (fig. S4. F and G) indeed revealed a strong increase in local curvatures of the lamina in cells with nuclear aggregates compared to control (~ 3-fold, n = 6 and 7 cells; fig. S4, H and I).

Using a total membrane stain (mCLING), we then visualized the NE in cells with or without nuclear aggregates. In control cells, the NE appeared as a continuous structure with minimal curvature, distinct from other cellular membranes. The presence of nuclear aggregates also increased local membrane curvature (Fig. 3G), resembling the increased lamin B1 curvatures found in expanded (fig. S4, D-G) and non-expanded (Fig. 2, G and J) cells. Compared to control, cells with nuclear aggregates had a ~2-fold increase in average NE curvature (n = 4 and 5 cells; Fig. 3, G-H; fig. S5, A and B). Together, these findings reveal that cells with nuclear aggregates display a strong deformation and disruption of their lamina and nuclear envelope (*24, 27*), which likely underlies the increased NE rupture frequencies in these cells (*19, 21*).

Our live-cell experiments revealed that, in the presence of nuclear aggregates, restoration of NE integrity occurred more slowly and was often completely absent, suggesting failed NE repair. To facilitate nanoscale observation of the NE at rupture sites, we used TREx on cells expressing nuclear aggregates and cGAS and stained with mCLING. Interestingly, we found multiple instances of rupture sites marked by intranuclear cGAS accumulation, where membrane deformations and disruptions were apparent around nuclear aggregates (Fig. 3, J and K; fig. S5, F and G). These observations suggest that polyQ fibrils often directly interfere with membrane resealing after NE rupture (*18*).

Huntington’s disease primarily affects neurons in various brain regions (*28–30*). Therefore, we expressed expanded or non-expanded polyQ protein in primary rat hippocampal neurons. Co-expression of mCherry-cGAS allowed for visualization of nuclear ruptures in these cells. As expected, expression of non-expanded, polyQ23-NLS protein in neurons did not result in aggregate formation (Fig. 4A), and only a small portion of polyQ23-NLS expressing cells showed cGAS accumulation in the nucleus (5.1±2.0%, n = 519 cells; Fig. 4B). Similar to expression in U2OS cells, neurons expressing polyQ74-NLS exclusively formed intranuclear aggregates (Fig. 4A). Interestingly, in contrast to the exclusively cytoplasmic localization in U2OS cells, expression of non-targeted polyQ74 in neurons resulted in the formation of both cytosolic and nuclear aggregates in the majority of cells (58.3±3.8%, n = 264 cells; Fig. 4A), resembling the localization of native non-targeted polyQ aggregates in HD (*29–31*). We found a strong increase in the amount of rupture events in neurons with polyQ74 (~4.0-fold, 20.6±5.8%, n = 264 cells) and polyQ74-NLS (~5.8-fold, 29.6±3.2%, n = 388 cells) aggregates, compared to polyQ23-NLS controls (Fig. 4B). Rupture events were sometimes accompanied by local disruptions in the nuclear lamin B1 signal (Fig. 4C), similar to disruptions in U2OS cells (Fig. 2, G and K). Together these findings demonstrate that nuclear aggregate-induced NE ruptures also occur in the cell type primarily affected in HD.

**Fig. 4.**
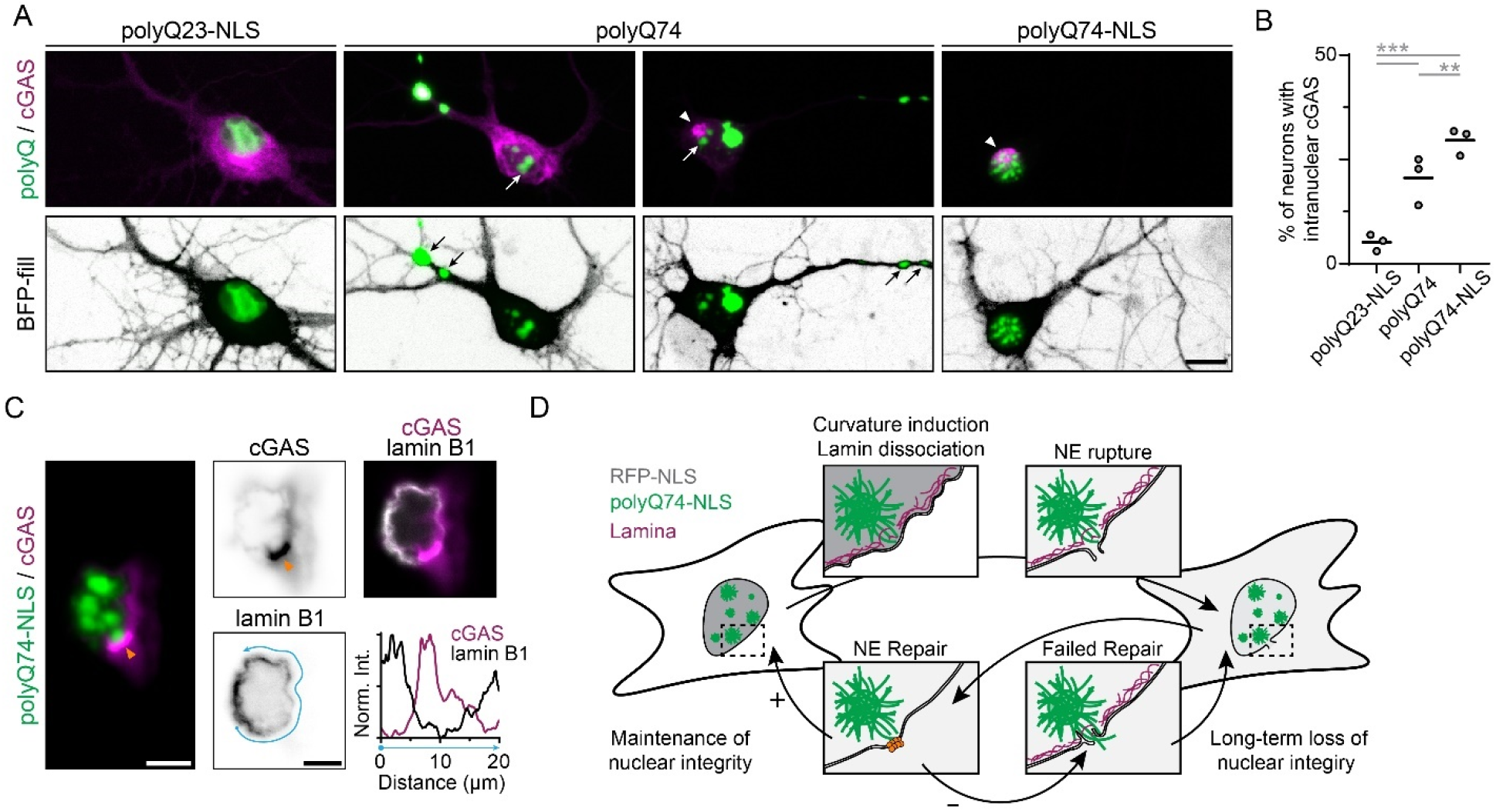
Nuclear polyQ aggregates induce NE rupture in primary rat hippocampal neurons. **(A)** Representative images of primary hippocampal neuron expressing BFP-fill (inverted greyscale), mCherry-cGAS (magenta) and polyQ23-NLS, polyQ74 or polyQ74-NLS (green). Neurons expressing polyQ74 show aggregation in soma, dendrites (black arrows) and nucleus (white arrows). Redistribution of mCherry-cGAS to the nucleus indicates NE rupture (white arrowhead). PolyQ signal was gamma adjusted (γ = 0.75). (**B**) Quantification of the percentage of neurons showing intranuclear cGAS accumulation (polyQ23-NLS, polyQ74, polyQ74-NLS; n=519, 264, 388; N=3). (**C**) Sum projections of confocal images of representative neuron showing lamin B1 (grey) depletion from rupture site indicated by mCherry-cGAS accumulation. (**D**) Proposed model for lamin disruption and sustained NE rupture induced by nuclear aggregates. Failure to restore nuclear integrity could lead to sustained loss of nuclear integrity. Scale bars indicate 10 μm (A) or 5 μm (C). **p<0.01, ***p<0.001.

Taken together, we have shown that while cytosolic and nuclear polyQ aggregates can both deform the nuclear lamina, only nuclear aggregates induce NE ruptures (Fig. 4D). Similar ruptures have been found in laminopathies and migrating cancer cells (*19, 32*), where they were reported to trigger nucleocytoplasmic mixing and DNA damage (*19, 21, 32, 33*). Crucially, while transient NE ruptures are often insufficient to trigger accumulation of endogenous cGAS and subsequent STING activation (*34, 35*), the prolonged loss of NE barrier function found in the presence of polyQ aggregates (Fig. 1F) could lead to a gradual buildup of detrimental effects, including nucleocytoplasmic mixing, transcriptional deregulation and even DNA damage, all features that are found in many HD models (*6–8*). Because several other neurodegenerative diseases also show intranuclear aggregation (*36–41*), alterations in nuclear morphology and impaired nuclear barrier function (*39, 41–49*), we speculate that nuclear aggregate-induced ruptures represent a unifying contributor to neurodegeneration that initiates a cascade of deregulated processes, culminating in degeneration and deleterious inflammation, a characteristic of most neurodegenerative diseases (*50, 51*).

## Acknowledgement

We thank Martin Hetzer for the gift of U2OS-RFP-NLS cells, Fulvio Reggiori and Anna Akhmanova for discussions and advice, Anne Janssen for early observations of nuclear ruptures in cells with aggregates, Hugo Damstra for help with expansion microscopy, and Klara Jansen for advice regarding primary neuron culture. This research was supported by the European Research Council (ERC Consolidator Grant 819219 to L.K.) and the Dutch Research Council (NWO, ZonMW 91217002 to H.H.K. and L.K.).

## Author Contributions

G.K. and L.C.K. designed the research. G.K., R.P. and B.H. created reagents and performed experiments. G.K. and R.P. analyzed the data and prepared figures. G.K. and L.C.K. wrote the manuscript with input from all coauthors. H.H.K. provided advice and guidance on aggregate biology and contributed reagents. L.C.K. supervised the study.

## Data and materials availability

All data are available in the main text or the supplementary materials. Plasmids are available from L.C.K. upon request.

## Supplementary Materials

Materials and Methods

Figs. S1 to S5

Movies S1 to S5

References (53-58)

## Supplementary figures and figure legends

**Fig. S1.**
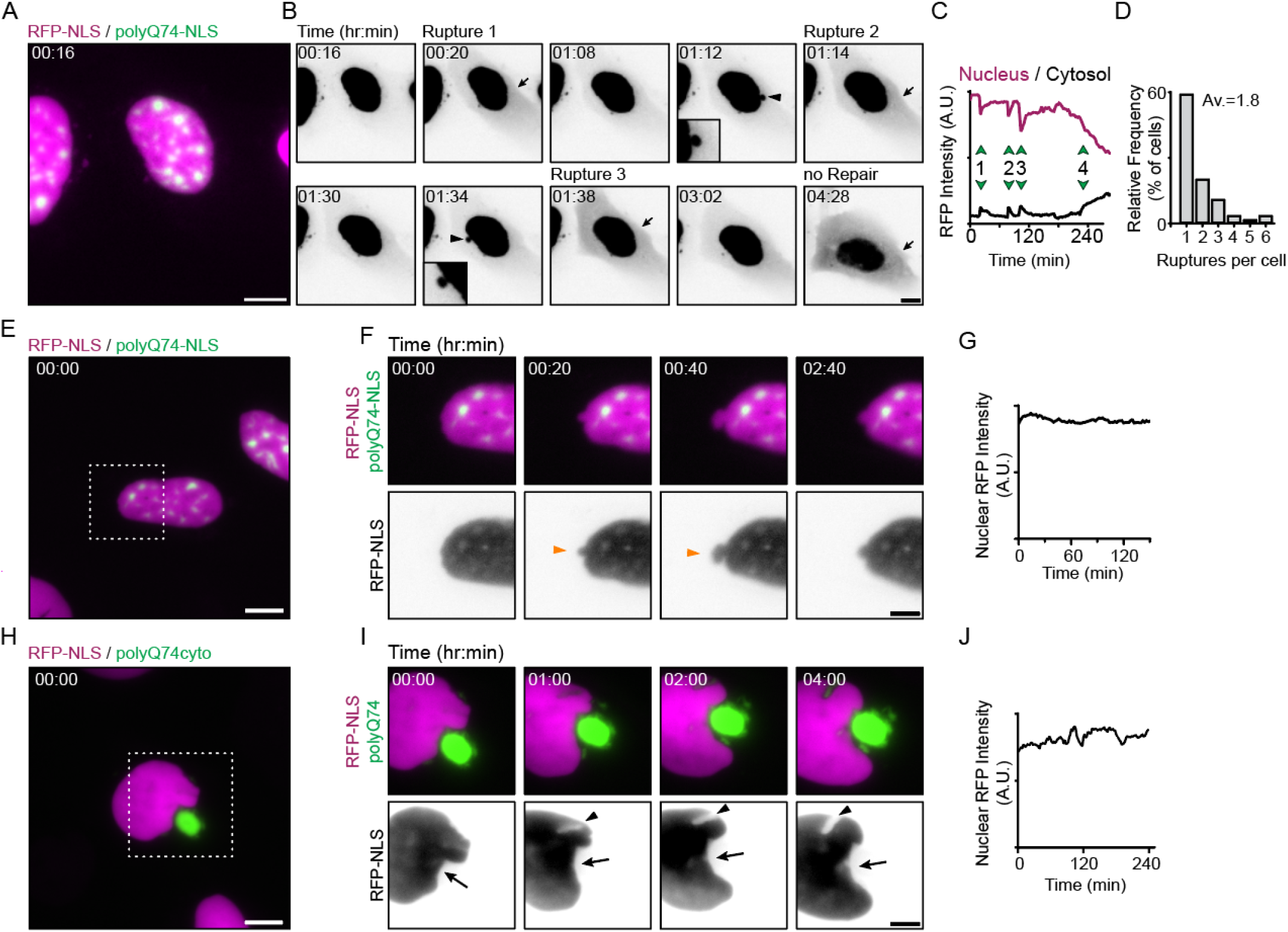
Studying NE rupture in cells expressing polyQ aggregates. (**A**) U2OS-RFP-NLS (magenta) cells expressing polyQ74-NLS aggregates (green). (**B**) Timelapse of RFP-NLS signal (inverted greyscale) of cell shown in (A) showing multiple blebbing events (black arrowheads) and NE rupture evidenced by cytosolic leaking of RFP-NLS signal (black arrow). (**C**) Graph showing nuclear (magenta curve) and cytosolic (black curve) RFP-intensity, indicating multiple instances of changes in nuclear and cytosolic RFP-NLS signal (green arrows 1, 2 and 3). After the last nuclear rupture (green arrow 4), nuclear enrichment does not restore. (**D**) Percentages of cells showing various amounts of nuclear ruptures during 8-hour imaging. (**E and F**) Timelapse images of representative U2OS-RFP-NLS cells (magenta and inverted greyscale) and polyQ74-NLS (green) showing blebbing event (orange arrowheads) without NE rupture. (**G**) Graph of nuclear RFP-intensity of timelapse in (F), showing no loss of RFP-NLS signal during blebbing event. (**H and I**) Timelapse images of representative U2OS-RFP-NLS cells expressing cytosolic polyQ74 aggregates (green), showing a characteristic bean shaped nucleus (black arrow) and NE deformation near a smaller cytoplasmic aggregate (black arrowhead). (**J**) Nuclear RFP-intensity of timelapse in (I) showing no loss of RFP-NLS signal during imaging. Scale bars indicate 10 μm (A, B, E and H) or 5 μm (F and I).

**Fig. S2.**
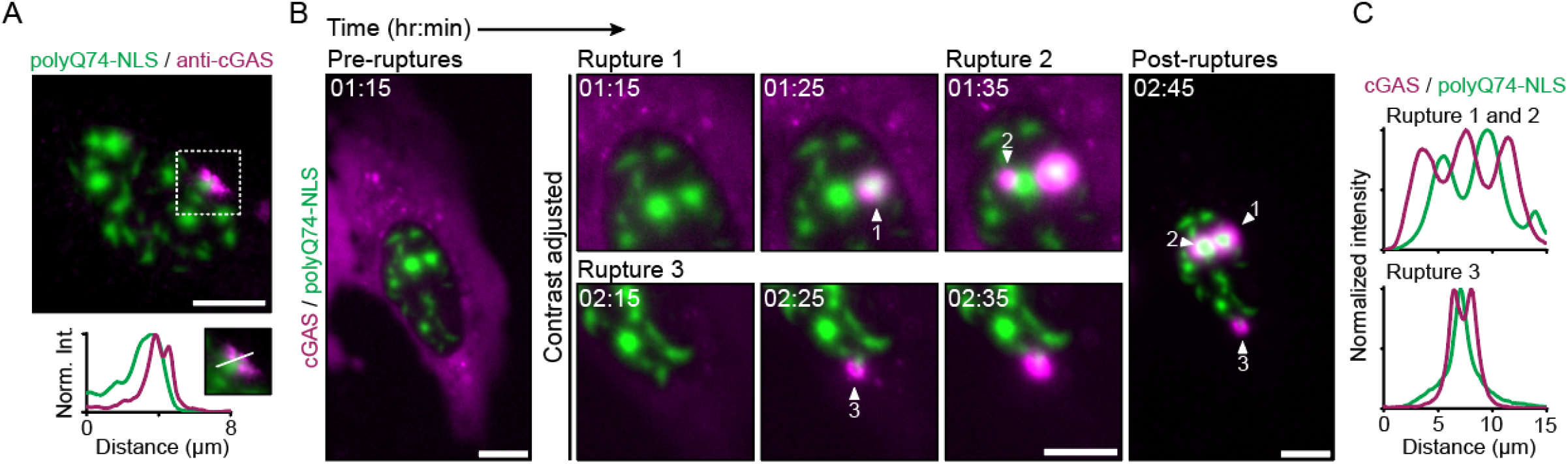
Accumulation of endogenous and overexpressed cGAS around polyQ aggregates. (**A**) Representative image of U2OSWT cells expressing polyQ74-NLS (green), immunostained for endogenous cGAS (magenta). Profile plot of line scan in inset, showing localization of endogenous cGAS around a nuclear aggregate. (**B**) Representative stills from live-cell imaging of U2OSWT cells expressing polyQ74-NLS (green) and mCherry-cGAS (magenta). Timelapse zooms show multiple ruptures (Rupture 1, 2 and 3) occurring at different aggregates. PolyQ and cGAS signal was gamma adjusted (γ = 0.75). (**C**) Graphs of intensity profiles across rupture sites shown in (B), indicating mCherry-cGAS accumulation around individual aggregates. All scalebars are 10 μm.

**Fig. S3.**
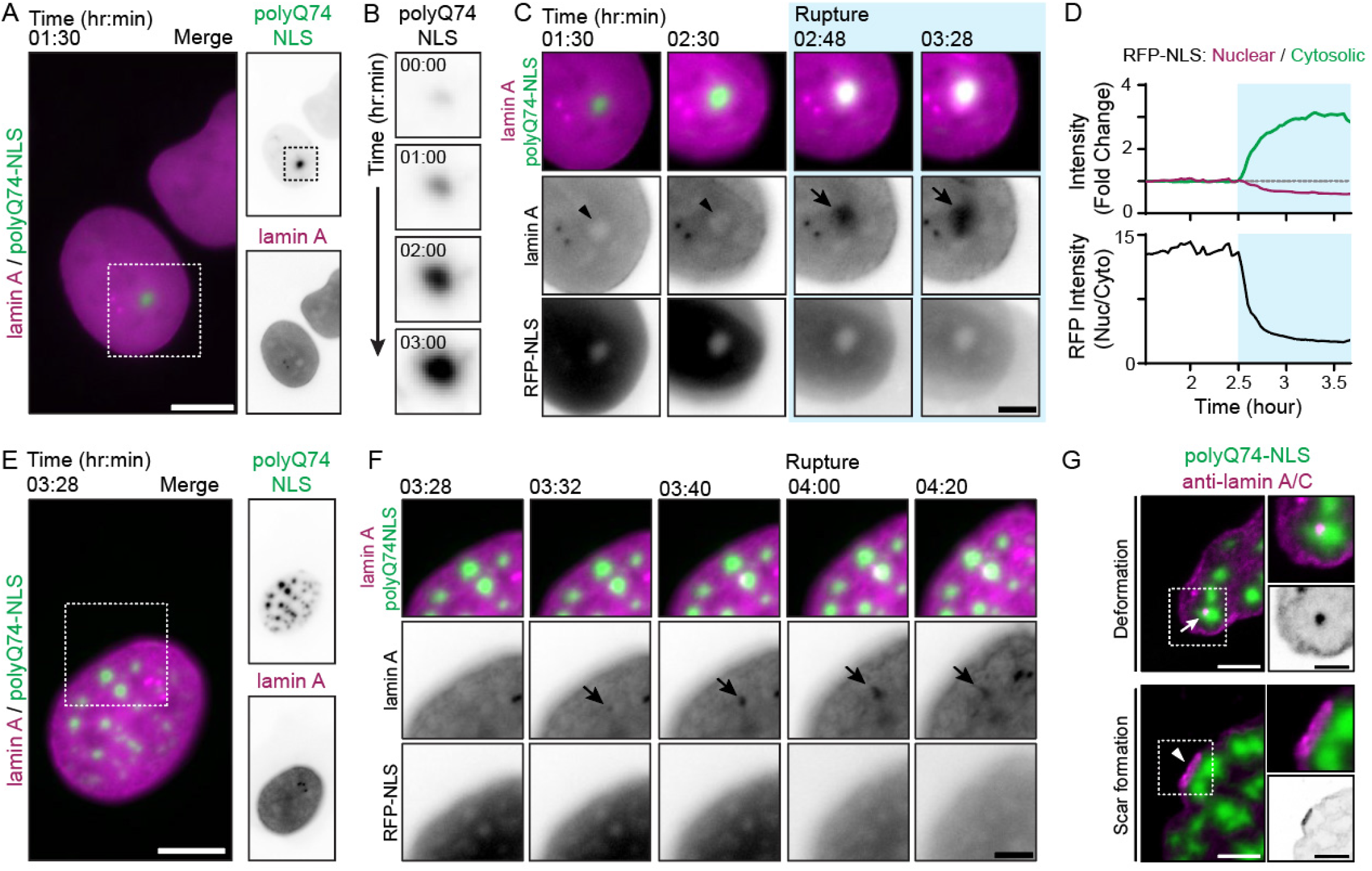
Lamina deformation and scar formation at nuclear aggregates. (**A and E**) Representative U2OS-RFP-NLS cells expressing polyQ74-NLS (green) and HaloTag-lamin A (magenta). (**B**) Timelapse zoom of cell shown in (A), showing aggregate growth during imaging. (**C**) Timelapse zooms of cell shown in (A). Stills show HaloTag-lamin A scar formation upon rupture (black arrowhead) at nuclear polyQ74-NLS aggregates. (**D**) Graph showing quantification of RFP-NLS intensity of stills shown in (C). Upper panel shows fold change in nuclear- (magenta curve) and cytosolic intensity (green curve). Bottom panel shows nuclear enrichment (nuclear/cytosolic intensity), indicating nuclear rupture during imaging. (**F**) Timelapse images of cell shown in (E) with minor HaloTag-lamin A accumulation (black arrow) at a nuclear aggregate. (**G**) Confocal microscopy images of representative U2OSWT cells expressing polyQ74-NLS aggregates (green). Minor deformation of endogenous lamin A/C (magenta, white arrow) and scar formation (white arrowhead). PolyQ signal (A, C, E and F) was gamma adjusted (γ = 0.75). Scale bar represents 10 μm (A and E), 5 μm (C, F and G) and 2.5 μm (G_zoom_).

**Fig. S4.**
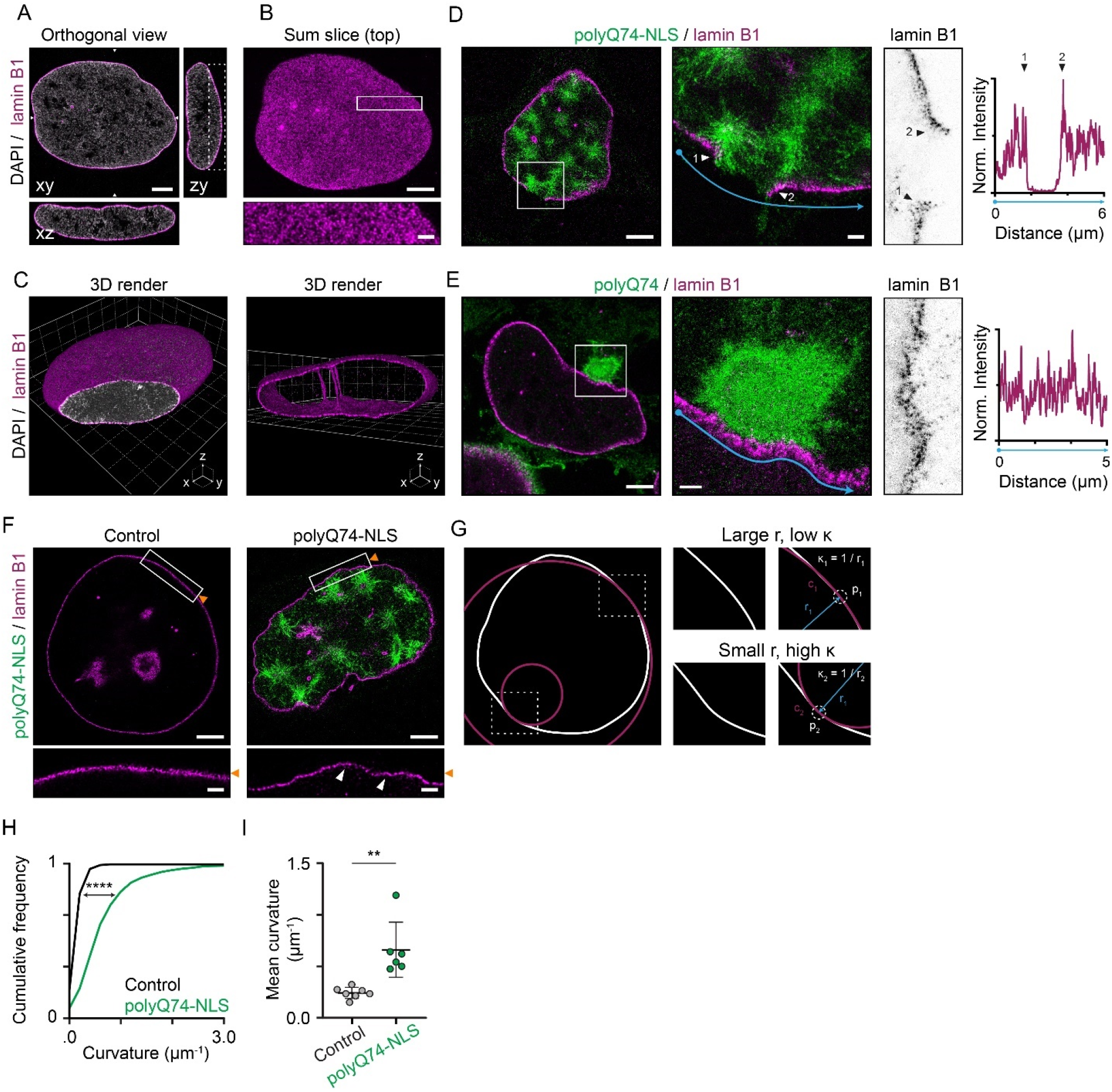
TREx microscopy reveals lamin B1 meshwork and deformations induced by nuclear polyQ aggregates. (**A-C**) Images of representative expanded U2OS cell immunolabelled for lamin B1 (magenta) and counterstained with DAPI (grey). (**A**) Orthogonal view showing discrete localization of lamin B1 at the nuclear periphery. (**B**) Sum projection of the top 1.5 μm of the nucleus shown in (A). Zoom shows lamin B1 meshwork. (**C**) 3D render of the whole lamin B1 meshwork of cell shown in (A). Cropped render of the middle of the nucleus reveals nucleoplasmic reticulum present in U2OS cells. (**D and E**) Representative images of expanded U2OS cells expressing polyQ74-NLS (D) or polyQ74 (E) stained for lamin B1. lamin B1 intensity profile indicates disrupted (E) or intact (E) lamin B1 meshwork at aggregate. (**F**) Sum projection of the middle plane of the nucleus of a representative control U2OS cells and a cell expressing polyQ74-NLS aggregates. Zooms show a piece of continuous lamina. White arrowheads indicate regions with high curvature in polyQ74-NLS expressing cells. (**G**) Schematic representation of segmentation of the lamin B1 meshwork from control cell shown in (F; white outline) and circles (magenta) fitted to this segmented meshwork to determine curvature. (**H and I**) Quantification of the distribution of lamina curvature (H) and mean curvature per cell (I) of control cells (black) and cells harboring polyQ74-NLS aggregates (green; 7 and 6 cells). Scale bars indicate 2.5 μm for overviews and 0.5 μm for zooms (A, B and D-F). Voxel sizes are 2 μm (grids in C). *p<0.05, ****p<0.0001.

**Fig. S5.**
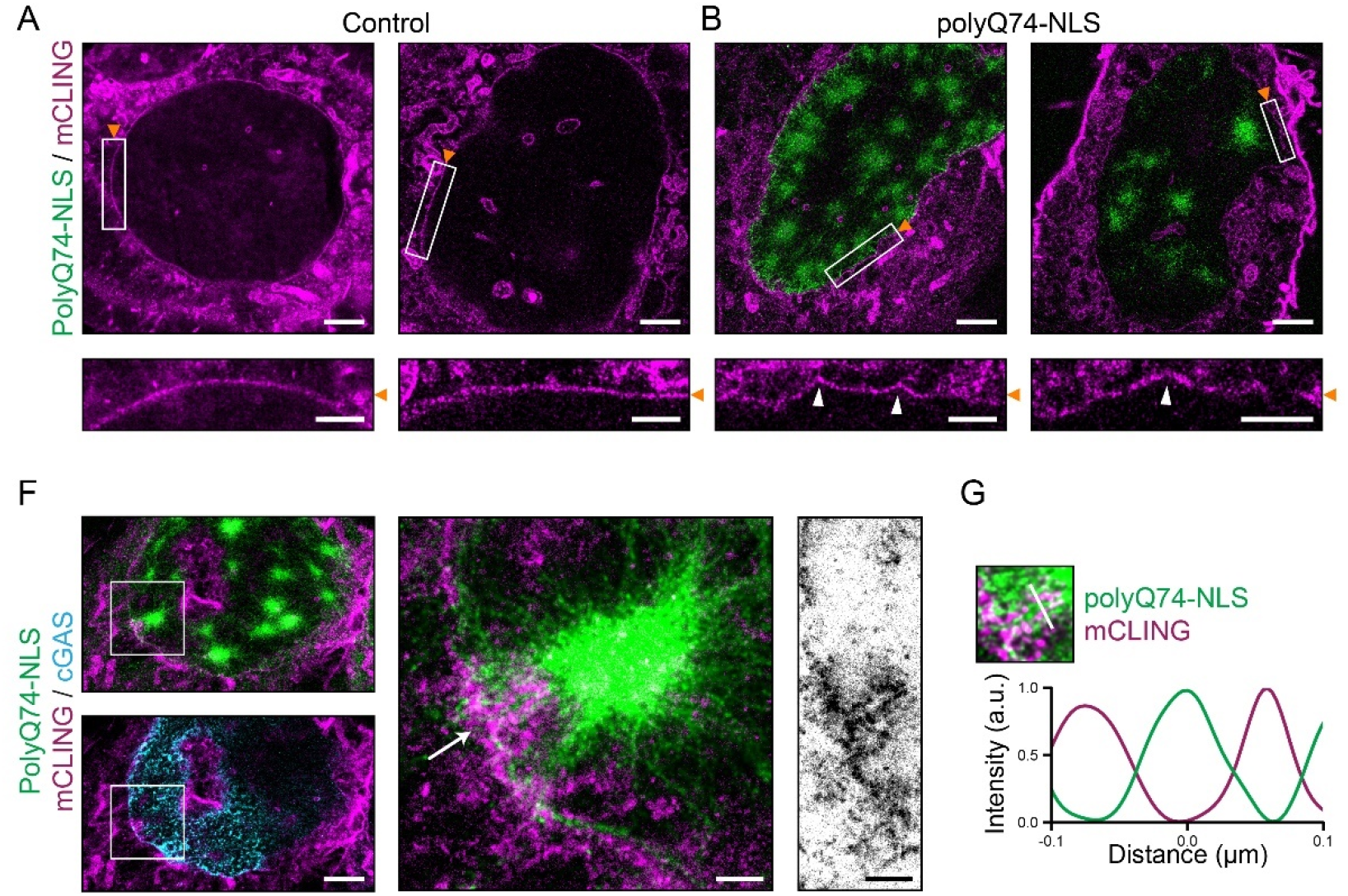
Deformation and disruption of the NE visualized using TREx microscopy. **(A-B)** Representative images of expanded U2OS control cells (A) or cells expressing polyQ74-NLS aggregates (green; B) stained for total membrane using mCLING (magenta). Zooms show a stretch of continuous nuclear membrane with high curvature (white arrowheads) in cells with nuclear aggregates. (**F**) Images of expanded U2OS cell expressing polyQ74-NLS and HA-mCherry-cGAS (cyan), showing NE deformation and invagination around a nuclear aggregate, at a NE rupture location marked by cGAS accumulation. (**G**) Intensity profile of NE rupture site shown in (F) showing mCLING stained NE accumulation around a polyQ fibril. Scale bars indicate 2.5 μm for overviews and 1 μm (A and B) or 0.5 μm (F) for zooms.

## Material and Methods

### Cell culture and Cell lines

U2OS-WT and U2OS-RFP-NLS cells were cultured in DMEM (Capricorn, HPSTA) supplemented with 10% FBS and 1% penicillin and streptomycin. Cells were kept at 37 °C and 5% CO_2_. U2OS-RFP-NLS cells were a gift from Martin Hetzer (Salk Institute, CA, USA) (*17, 20*) and were cultured in full DMEM supplemented with 0.5 mg/ml G418 (Abcam, ref. ab144261, lot. GR 162868-8). Cells were regularly tested for mycroplasma using MycoAlert Mycoplasma Detection Kit (Lonza).

### Constructs

The HaloTag-lamin A construct was generated by first inserting GFP-lamin A from pBABE-puro-GFP-lamin A into pEGFP-C2 using NheI and BamHI. Then, the HaloTag fragment was generated by PCR and inserted into pEGFP-lamin A using restriction digestion with NheI and BglII to generate HaloTag-lamin A. Constructs encoding for nuclear targeted or untargeted huntingtin exon1 fragment with poly-glutamine stretch (polyQ23-NLS, polyQ74-NLS and polyQ74cyto) were a generated previously (*16*). The mCherry-cGAS was a gift from Dennis E. Discher (UPENN, PA, USA) (*21, 52*). For visualization of rupture sites in TREx microscopy, mCherry-cGAS was inserted into an HA-containing vector (gift from Qing Zhong; Addgene plasmid #280274) (*53*) generating HA-mCherry-cGAS. pBABE-puro-GFP-lamin A was a gift from Tom Misteli (Addgene plasmid #17662) (*54*).

### Plasmids and transfection

U2OS WT and U2OS-RFP-NLS were plated on 23-mm or 18-mm diameter coverslips 1-2 days before transfection. One day before live-cell imaging or fixation, cells were transfected using FuGENE6 (Promega) at a ratio of 3 μl transfection reagent per 1 μg DNA. Cells were either fixed or used for live cell imaging 1 day after transfection.

### Rat Hippocampal Neuron Culture and Transfection

Primary hippocampal cultures were isolated from embryonic rat brain (day 18) as described previously (*55*). Cells were plated on coverslips coated with laminin (2μg ml^-1^) and poly-L-lysine (30μg ml^-1^). Cultures were grown in full Neurobasal medium (NB, Gibco, ref. 21103049) with B27 (Gibco, ref. 17504044), 0.5 mM glutamine, 12.5 μM glutamate and penicillin/streptomycin. Neurons were cultured at 37°C in 5% CO_2_ for 9 days prior to transfection. Per well, transfection was performed using a transfection mix containing 1.8 μg DNA and 3.3 μl lipofectamine 2000 (Invitrogen, ref. 11668019) in 200 μl NB. The transfection mix was thoroughly mixed and incubated for 30 min. Neurons were transfected by adding the transfection mix to neurons in NB with 0.5 mM glutamine for 60 minutes. During transfection neurons were kept at 37°C in 5% CO_2_. After transfection, neurons were washed with NB and returned to full NB. Neurons were fixed using 4% paraformaldehyde (PFA) with sucrose 48h after transfection.

### Live-cell and Fluorescence Microscopy

For live-cell imaging, cells were imaged on a Nikon Eclipse Ti with equipped with an incubator chamber (Tokai Hit; INUG2-ZILCS0H2) on a motorized stage (ASI). Illumination was performed using a CoolLED pE4000 (CoolLED) and ET-EGFP (49002, chroma), ET-mCherry (49008, chroma) and ET-CY5 (49006, chroma) filters. All images were acquired with a Coolsnap HQ2 CCD camera (Photometrics). The microscope was controlled using μManager software (*56*). Cells were imaged in a metal imaging chamber (Invitrogen, ref. A7816) that was sealed by placing a coverslip on top to prevent medium evaporation.

For long-term timelapse imaging used for quantification of nuclear envelope rupture, blebbing and recovery in U2OS-RFP-NLS, cells were imaged using a 20X dry objective (Plan Apo. NA 0.75, Nikon) every 5 minutes for 8 hours. All other live cell imaging was done using a 40X oil immersion objective (Plan Fluor, NA 1.3, Nikon). Cells were imaged every 2-3 minutes for 2-6 hours. For experiments with HaloTag-lamin A, cells were incubated with JF 646 (Janelia Fluor, ref. GA112A, lot. 0000486504) for 30 min prior to imaging. For quantification of NE ruptures, cells were treated with 2mM thymidine (Calbiochem, ref. 6060-5GM, lot. D0017544) to prevent NE breakdown caused by mitosis.

Fixed-cell immunofluorescence microscopy images used for scoring of lamin B1 phenotype or cGAS intranuclear accumulation were taken on a Nikon Eclipse Ni-U microscope with a 100x oil immersion objective (plan Apo Lambda, N.A. 1.44, Nikon) equipped with ET-EGFP (49002, Chroma) and ET-mCherry (49008) filters. All other fixed cell imaging was performed using a point-scanning confocal Zeiss AiryScan LSM880 microscope using a 63x immersion objective (Plan-Apochromat, 1.2 NA), controlled by Zen Black software.

Images of the expanded cells were acquired using a Leica TCS SP8 STED 3X microscope equipped with an HC PL APO ×86/1.20W motCORR STED (Leica 15506333) water objective controlled using Leica Application Suite X.

### Antibodies and Reagents

For immunofluorescence labeling the following antibodies were used; anti-lamin A/C (Santa Cruz, ref. sc-7292, lot. L1919), anti-lamin B1 (Abcam, ref. ab160848, lot. GR3417466-1), anti-cGAS (Cell Signaling Technology, ref. 15102, lot. 4), anti-GFP (MBL-Sanbio, ref. 598, lot. 081), anti-HA (Santa Cruz, ref. SC-57592, lot. L2310) and anti-GFP (Aves Labs, ref. GFP-1010, lot. GFP3717982). Secondary antibody labelling was done using goat anti-rabbit 488 (Thermo Fischer Scientific, ref. A11034, lot. 2256692 and 2286890), goat anti-rabbit 568 (Thermo Fischer Scientific, ref. A11036, lot. 2045347), goat anti-mouse 568 (Thermo Fischer Scientific, ref. A11031, lot. 2124366), goat anti-mouse 594 (Thermo Fischer Scientific, ref. A11032, lot. 2069816 and 2397936), goat anti-chicken 488 (Thermo Fischer Scientific, ref. SA5-10070, lot. VI3075603) and goat anti-rabbit 594 (Thermo Fischer Scientific, ref. A11037, lot. 2160431).

### Immunofluorescence

For immunofluorescence labelling for wide-field or confocal fixed-cell imaging, cells were first fixed using pre-warmed 4% PFA. Cells were washed with 1X PBS (Lonza), permeabilized using 0.2% Triton-X100 and blocked using blocking solution (3% BSA in 1X PBS) for 1 hour at RT. Cells were incubated with primary antibody (1:500 in blocking solution) at 4°C overnight. Cells were subsequently washed in 1X PBS and incubated with the appropriate secondary antibodies (1:500 in blocking solution) for 1hr at RT. Cells were dried and mounted using Prolong Diamond Antifade Mountant (Invitrogen, ref. P36965).

### Expansion microscopy

#### Fixation and pre-extraction

For tenfold robust expansion (TREx) microscopy, we adapted the protocol described previously (*57*). For mCLING staining, cells were fixed using pre-warmed 4% PFA, 4% sucrose (w/v) and 0.1% glutaraldehyde. Coverslips were incubated with 10 μM mCLING (Synaptic Systems, 710 006AT1) for 4 to 6h at 37 °C and subsequently incubated overnight at RT. The incubated samples were post-fixed with pre-warmed 4% PFA and 0.1% glutaraldehyde. Cells that were not stained using mCLING were only fixed with pre-warmed 4% PFA. Samples that were pre-extracted were incubated with 1 ml 0.15% (v/v) pre-warmed TritonX-100 in PBS for 1 minute prior to fixation.

#### Gelation and expansion

The following monomer solution was prepared on ice: 1.085 M sodium acrylate (Sigma-Aldrich, 408220), 2.664 M acrylamide (Sigma-Aldrich, A4058) and 0.009% (v/v) N,N’-methylenebisacrylamide (Sigma-Aldrich, M1533) in 1x PBS. The polymerization reaction was started by addition of 1.5% (v/v) tetramethylethylenediamine (TEMED) and 1.5% (v/v) ammonium persulfate (APS). The monomer solution was vortexed and 170 μL (per coverslip) was pipetted into a silicon gelation chamber attached to a parafilm covered glass slide. The coverslip containing stained cells was blotted onto the monomer solution. The gelation chambers were incubated at 37°C for one hour. Subsequently, gels were digested in 2 ml digestion mix for 4h at 37 °C and expanded up to 10x using MilliQ.

#### Data processing and 3D-rendering

Prior to image analysis, all TREx images except mCLING channels were deconvolved with Huygens Professional version 21.04 (Scientific Volume Imaging, The Netherlands, http://svi.nl) using the CMLE algorithm with 4 SNR and 20 iterations. mCLING channels were blurred using Gaussian Blur 3D with 0.8 sigma (both X, Y, and Z) and subsequently used to generate a rolling average using Running Z projector (https://valelab4.ucsf.edu/~nstuurman/IJplugins/Running_ZProjector.html) with a running average size of 3 slices per slice. For 3D volume rendering Arivis Vision4D version 3.5.0 was used. PolyQ74-NLS filaments were processed by normalizing the intensity (method: simple), detected using a “random forest” machine learning classifier (https://ukoethe.github.io/vigra/), segmented by intensity thresholding, and filtered by size (> 0.001 um^3^) and manual deletion. The lamin B1 network was processed by normalizing the intensity (method: simple) and masking external nuclear signal using a manually generated mask.

### Quantifications

Cells were manually scored for quantification of at least one nuclear envelope rupture or nuclear blebbing event during 8-hour imaging. Cells that divided, died or migrated out of the field of view of the camera were excluded from the analysis. Cells that did not have nuclear RFP-NLS enrichment at the start of imaging were also excluded from the analysis. The fraction of cells that recovered nuclear enrichment of RFP-NLS intensity were determined by manually scoring recovery in all cells that showed a nuclear rupture event. Cells that ruptured in the last hour of imaging (7-8 hours after start of imaging) were excluded from the analysis. For quantification of recovery dynamics after nuclear rupture, RFP-NLS intensity ratios (Nuclear/Cytoplasmic) were normalized to the average intensity ratio of 3 frames before rupture.

Distance between a rupture site indicated by intranuclear cGAS accumulation and nuclear polyQ74-NLS aggregates was determined by placing a ROI around the cGAS accumulation and nearest polyQ74-NLS aggregate on the first frame after rupture. The centers of these ROIs were calculated using Imagej/Fiji and the distance between these centers was taken as the distance between the rupture location and polyQ74-NLS aggregate.

The effect of polyQ aggregate presence in the nucleus or cytosol on the nuclear lamina was determined by manually scoring endogenous lamin B1 phenotypes in control U2OSWT cells or cells expressing polyQ23-NLS, polyQ74-NLS or polyQ74. The percentage of cells showing either lamin B1 disruption (absence of lamin B1 signal at nuclear rim), multifocal deformation (wrinkling of large part of lamin B1 signal at nuclear rim) or unifocal deformation (single deformation of lamin B1; Fig. 2H).

Nuclear rupture frequency in neurons was done by manually scoring intranuclear cGAS accumulation in cells that were transfected (control and polyQ23-NLS) or showed aggregates (polyQ74-NLS and polyQ74). Neuron viability was assessed by neuronal morphology in BFP-fill signal. Neurons that were dead or that did not express sufficient BFP-fill to visualize neuronal processes were excluded from the analysis.

Nuclear curvature was calculated in ImageJ/Fiji (https://gist.github.com/lacan/42f4abe856f697e664d1062c200fd21f). The nuclear outline was traced manually on a slice with a well-defined nuclear membrane based on the mCLING signal and smoothened using the fit spline function. For each vertex point on the outline, a circle was fitted on the outline. The radius of this circle was used to calculate the curvature (κ = 1/r; fig. S4, E).

### Statistical Analysis

Prism9 (GraphPad) was used for generating all graphs and statistical analyses. Differences in nuclear rupture frequency, blebbing frequency and recovery frequency were tested for statistical significance using a Fischer’s exact test with the null hypothesis that proportions were equal in all groups. Differences in percentages of hippocampal neurons showing intranuclear cGAS accumulation were also tested for statistical significance using Fischer’s exact test. Lamin B1 phenotypes were tested for significant differences using an ordinary one-way ANOVA and Tukey’s post hoc test or Fischer’s exact test. All Fischer’s exact tests were corrected for multiple testing using Bonferroni correction.

Recovery dynamics after NE rupture were measured by first calculating traces of nuclear enrichment (Nuclear enrichment = I_nuclear RFP_/I_cytosolic RFP_) per cell normalized to pre-rupture enrichment values. Average nuclear enrichment was calculated by aligning normalized enrichment values to the time of NE rupture.

To test whether average NE curvature of control cells was significantly different from cells harboring polyQ74-NLS aggregates we used an unpaired Student’s t-test. We also determined whether the cumulative frequency distributions of NE curvature were different between these groups using a Mann-Whitney test.

All quantifications of intensity values in U2OSWT and U2OS-RFP-NLS cells expressing polyQ74, polyQ74-NLS, mCherry-cGAS or HaloTag-lamin A were performed on raw, unprocessed, images. Values were background-corrected before plotting.

### Data Processing

All image processing of non-expanded samples was performed using FIJI (*58*). Multiple images of intense signal of polyQ-aggregates or cGAS were gamma corrected (γ = 0.50-0.75) to increase visibility of signal around intense cores (see figure legend). Final figure panels were prepared in Adobe Illustrator.

